# Barley RIC157 is involved in RACB-mediated susceptibility to powdery mildew

**DOI:** 10.1101/848226

**Authors:** Stefan Engelhardt, Adriana Trutzenberg, Katja Probst, Johanna Hofer, Christopher McCollum, Michaela Kopischke, Ralph Hückelhoven

**Affiliations:** Phytopathology, TUM School of Life Science Weihenstephan, Technical University of Munich, Emil-Ramann-Str.2, 85354 Freising, Germany

## Abstract

Successful obligate pathogens benefit from host cellular processes. For the biotrophic ascomycete fungus *Blumeria graminis* f.sp. *hordei* (*Bgh*) it has been shown that barley RACB, a small monomeric G-protein (ROP, RHO of plants), is required for full susceptibility to fungal penetration. The susceptibility function of RACB probably lies in its role in cell polarisation, which may be co-opted by the pathogen for invasive ingrowth of its haustorium. However, the actual mechanism of how RACB supports the fungal penetration success is little understood. RIC proteins (ROP-Interactive and CRIB-(Cdc42/Rac Interactive Binding) motif-containing) are considered scaffold proteins which can interact directly with ROPs via a conserved CRIB motif. Here we describe a yet uncharacterised RIC protein, RIC157, which can interact directly with RACB *in planta*. We show that RIC157 undergoes a recruitment from the cytoplasm to the cell periphery in the presence of activated RACB. During fungal infection, RIC157 and activated RACB colocalise at the penetration site, particularly at the haustorial neck. In a RACB-dependent manner, transiently overexpressed RIC157 renders barley epidermal cells more susceptible to fungal penetration. This suggests that RIC157 promotes fungal penetration into barley epidermal cells via its function downstream of RACB.

## Introduction

Plants have developed a multilayered immunity to defend microbial invasion. This consists of pre-formed barriers and induced defences that base on the receptor-mediated recognition of microbe-derived and endogenous elicitors (Boller and Felix 2009; Stukenbrock and McDonald 2009). Except for necrotrophs, invading microbes rely to different extents on a living host to establish an infection. With the help of secreted effectors, pathogens undermine plant immune reactions and influence the host metabolism to render their micro-environment more favourable (Białas *et al.*, 2018, Han and Kahmann 2019). The co-evolution between microbial effectors and their specific host target molecules can lead to an increase in host specialisation, symbiotic relationships thereby demonstrating extreme examples. Regarding pathogens, especially in genomes of cereal powdery mildew fungi it has been observed that the amount of genes encoding for metabolic enzymes is massively reduced, concomitantly with the proliferation of the putative effector gene pool and transposable elements (Spanu *et al.*, 2010, Wicker *et al.*, 2013, Frantzeskakis *et al.*, 2018). Plant targets of these effectors are not necessarily involved in resistance mechanisms, but also in cellular processes that, when controlled by the pathogen, can support the susceptibility towards the invading pathogen. With the current possibilities to use a plethora of different breeding technologies, durable crop resistance based on the loss of these susceptibility gene product functions is within reach (Dangl *et al.* 2013, Engelhardt *et al.*, 2018).

Powdery mildew fungi infect a huge variety of monocot and dicot plants causing massive yield losses in crops. The ascomycete fungus *Blumeria graminis* f.sp. *hordei* (*Bgh*) is the specific causal agent of the agronomically important powdery mildew disease on barley (*Hordeum vulgare*) (Jørgensen and Wolfe 1994). As an obligate biotrophic parasite, *Bgh* requires living epidermal cells to complete its life cycle. Airborne conidia germinate on the leaf surface and form an appressorium to penetrate the cuticle and the cell wall with the help of an immense turgor pressure and the release of cell wall-degrading enzymes (McKeen and Rimmer 1973; Schulze-Lefert and Vogel 2000; Hückelhoven and Panstruga 2011). A successful fungal infection is characterised by the formation of a haustorium inside the host cell, which is essential for nutrient uptake and effector protein delivery (Hahn and Mendgen 2001, Voegele *et al.*, 2001, Panstruga and Dodds 2009). The haustorium is separated from the host cytosol by the extrahaustorial matrix and surrounded by the extrahaustorial membrane (EHM), which is continous with the plant plasma membrane, but differs functionally and biochemically from it (Koh *et al.*, 2005, Inada and Ueda 2014, Kwaaitaal *et al.*, 2017). It is feasible to imagine a pathogen-triggered active contribution of the plant to accomodate the fungal haustorium.

ROPs (RHO (RAS homologue) of plants, or RACs, for rat sarcoma (RAS)-related C3 botulinum toxin substrate) form a unique subfamily of small monomeric RHO GTPases in plants, since they do not fall into the phylogenetic RHO subclades of RAC, CDC42 and RHO GTPases found in yeast or animals (Brembu *et al.*, 2006). G-proteins are paradigms of molecular switches due to their ability to bind and hydrolyze GTP. The GTP-bound form represents the activated state, and a plasma membrane association of ROP-GTP via posttranslational lipid modifications is required for downstream signalling (Yalovsky 2015). Upon GTP hydrolysis, GDP-bound or nucleotide-free ROPs are inactive in downstream signalling. The cycling between activated and inactive state needs to be spatiotemporally controlled by regulatory partners. Guanine nucleotide exchange factors (GEFs) positively regulate ROP activity by facilitating the GDP/GTP exchange. In plants, three different sorts of ROP GEFs can be distinguished based on their particular GEF domain: PRONE (plant-specific Rop nucleotide exchanger), DHR2 (DOCK homology region 2, found in SPIKE1) and a less well characterized DH-PH domain (B-cell lymphoma homology-pleckstrin homology) described in a plant homolog of human SWAP70 (Berken *et al.*, 2005, Meller *et al.*, 2005, Gu *et al.*, 2006, Basu *et al.*, 2008, Yamaguchi and Kawasaki 2012, Yamaguchi *et al.*, 2012, He *et al.*, 2018). The interaction of ROPs with a GTPase Activating Protein (GAP) enhances the intrinsic GTP hydrolysis activity, followed by ROP inactivation (Berken and Wittinghofer 2008). Beside their putative involvement in ROP recycling, Guanine nucleotide Dissociation Inhibitors (GDIs) bind and sequester inactive ROPs in the cytoplasm and are therefore considered negative regulators of ROP activity (Klahre *et al.*, 2006, Boulter and Garcia-Mata 2010). ROPs are involved in the regulation of a multitude of cellular processes. For instance, the cytoskeleton organisation and consequentially cell shape and function is subject to RHO-like GTPase control (Chen and Friml 2014). In *Arabidopsis thaliana* xylem vessels, AtROP11 signaling promotes cell wall apposition and shapes cell wall pit boundaries (Sugiyama *et al.*, 2019). Different ROPs are involved in polar cell growth and even function antagonistically during the generation of *Arabidopsis thaliana* pavement cells (Craddock *et al.*, 2012). Beside cell polarisation and cytoskeleton organisation, ROPs have been also implicated in membrane trafficking and auxin signaling (Yalovsky *et al.*, 2008, Wu *et al.*, 2011). OsRac1 from rice (*Oryza sativa* enhances cell division by regulating OsMAPK6, thereby promoting rice grain yield (Zhang *et al.*, 2019). OsRAC1 has also been demonstrated to regulate immune-related processes like ROS production, defense gene expression and cell death. OsRac1 becomes activated by OsRacGEF1 upon receptor-mediated perception of fungal-derived chitin by OsCEBiP and OsCERK1 (Akamatsu *et al.*, 2013). Chitin-perception might also lead to the activation of OsRAC1 by OsSWAP70 (Yamaguchi *et al.*, 2012). Downstream signaling by OsRAC1 is also triggered after recogniton of pathogen effector proteins: Plasma membrane-localised Pit, a nucleotide binding-leucine rich repeat resistance (NLR) protein for the rice blast fungus *Magnaporte oryzae*, associates with DOCK family GEF OsSPK1, thereby likely activating OsRac1 (Kawano *et al.*, 2010, Kawano *et al.*, 2014, Wang *et al.*, 2018). A recent report regarding an involvement in defence reactions against rice blast mediated by the NLR protein PID3 (Zhou *et al.*, 2019) opens up the possibility of OsRac1 being a downstream hub of other rice NLR proteins.

In the barley-powdery mildew interaction, several barley proteins involved in ROP signaling or ROP activity regulation have been shown to influence fungal penetration success. The barley ROP RACB has been shown to act as susceptibility factor (Schultheiss *et al.*, 2002, Schultheiss *et al.*, 2003, Hoefle *et al.*, 2011). In the absence of the pathogen, RACB appears to be involved in cell polarization processes, as stable RACB silencing affects stomatal subsidiary cell and root hair development (Scheler *et al.*, 2016). The expression of a constitutively activated GTP-bound RACB supported fungal penetration success into barley epidermal cells, whereas silencing RACB by RNA interference (RNAi) renders epidermal cells less susceptible to fungal invasion. Two RACB-interacting proteins have been described as negative regulators of RACB function in susceptibility. First, the Microtubule-Associated ROP-GAP1 (MAGAP1) is recruited to the cell periphery by activated RACB and limits susceptibility to powdery mildew likely by enhancing the GTP-hydrolizing activity of RACB (Hoefle *et al.*, 2011). Second, activated RACB interacts with the cytoplasmic ROP binding kinase1 (RBK1) *in vivo* and enhances its kinase activity *in vitro* (Huesmann *et al.*, 2012). Transient silencing of RBK1 or RBK1-interacting protein SKP1 (type II S-phase kinase1-associated protein) suggested that RBK1 acts in negative regulation of RACB protein stability and hence in disease resistance (Reiner *et al.*, 2016).

In order to regulate cellular processes, ROPs need to activate or deactivate downstream executors (otherwise called ROP effectors, which is avoided here to distinguish from pathogen effectors). The interaction to some of these executors is often indirect and achieved via scaffold proteins bridging the activated ROPs to their signal destination targets. Some ROP scaffold proteins have been described so far in detail, RACK1, ICR/RIPs and RICs. Rice RACK1 (Receptor for Activated C-Kinase 1) interacts with several proteins in the OsRac1 immune complex supporting a role in rice innate immunity (Nakashima *et al.*, 2008). ICR/RIPs (Interactor of Constitutive Active ROP/ROP Interactive Partners) are required for cell polarity, vesicle trafficking and polar auxin transport (Lavy *et al.*, 2007, Hazak *et al.*, 2014). In barley, RIPa interacts with RAC1 and organizes microtubule arrays in concert with MAGAP1 (Hoefle *et al.*, 2020). Barley RIPb interacts directly with RACB and enhances disease susceptibility towards powdery mildew (McCollum *et al.*, 2019 Preprint). RIC (ROP-Interactive and CRIB-domain containing) proteins, another class of scaffold proteins in ROP signaling, share a highly conserved CRIB motif (Cdc42/Rac Interactive Binding motif, Burbelo *et al.*, 1995), which is essential for the direct interaction with ROPs (Wu *et al.*, 2001). The CRIB domain is also present in a subset of ROP GAPs such as barley MAGAP1 (Schaefer *et al.*, 2011; Hoefle *et al.*, 2011). In barley, the knowledge about RIC protein functions is quite limited. RIC171, however, has been shown to not only interact directly with RACB, but also to increase fungal penetration efficiency in barley epidermal cells upon overexpression. Activated RACB recruits RIC171 to the cell periphery and, in the presence of *Bgh*, RIC171 accumulated at the haustorial neck close to the penetration site (Schultheiss *et al.*, 2008). In *Arabidopsis thaliana* (At), 11 different RIC proteins have been identified, that do not share common sequence homology outside their CRIB domain (Wu *et al.*, 2001). By directly interacting with AtROPs, AtRIC proteins are involved in numerous cellular processes. During salt stress, AtROP2 regulates microtubule organisation in an AtRIC1-dependent manner (Li *et al.*, 2017). AtRIC1 also interacts with AtROP6 in pavement cells to enhance the ordering of cortical microtubules upon hormonal signals (Fu *et al.*, 2009) and is involved in cell elongation during pavement cell morphogenesis (Higaki *et al.*, 2017). AtRICs counteract each other to a certain extent as well, as seen with AtROP1-interacting AtRIC3 and AtRIC4 during pollen tube growth. AtRIC3 regulates calcium influx and triggers actin depolymerisation, whereas AtRIC4 enhances actin polymerisation (Gu *et al.*, 2005). Light-induced stomatal opening is regulated via the AtROP2-AtRIC7 pathway. AtROP2 and AtRIC7 are likely to impinge on vesicular trafficking by inhibiting AtExo70B1, which results in a diminished stomatal opening (Hong *et al.*, 2016). These examples emphasize the importance of ROP proteins as signaling hubs for various developmental processes as well as the role of RIC proteins in finetuning specific cellular responses.

Here we show results on barley RIC157, a CRIB domain-containing protein that interacts CRIB motif-dependently with RACB in yeast and *in planta*. Overexpression of RIC157 increases the powdery mildew penetration efficiency in barley leaf epidermal cells in a RACB-dependent manner. Cytosolic RIC157 is recruited to the cell periphery specifically by activated RACB and both proteins co-localise at the haustorial neck during the compatible interaction with *Bgh*. Our findings indicate a possible role of the RACB-RIC157 signaling module in promoting fungal penetration, thereby increasing susceptibility towards *Bgh*.

## Results

### Identification of RIC proteins in barley

Except for a couple of amino acids, individual RIC proteins typically show a lack of primary sequence homology to other proteins in the database outside their CRIB domain (Wu *et al.*, 2001). The highly conserved CRIB motif has been shown to interact directly with activated small RHO GTPases (Burbelo *et al.*, 1995, Aspenström 1999). In order to identify additional RIC proteins in barley, we performed a BLAST search using the CRIB motif of previously described RACB interacting protein RIC171 (Schultheiss *et al.*, 2008) against the 2019 annotation of all barley coding sequences (Barley all CDS Morex v2.0 2019, https://webblast.ipk-gatersleben.de/barley_ibsc/). Beside RIC171, we identified another seven proteins sharing the properties of RIC proteins, and named them according to their predicted amino acid sequence length RIC153 (HORVU.MOREX.r2.3HG0258770), RIC157 (HORVU.MOREX.r2.6HG0469110), RIC163 (HORVU.MOREX.r2.5HG0443720), RIC168 (HORVU.MOREX.r2.2HG0170820), RIC170 (HORVU.MOREX.r2.6HG0521090), RIC171 (HORVU.MOREX.r2.2HG0164690), RIC194 (HORVU.MOREX.r2.3HG0258620), RIC236 (HORVU.MOREX.r2.2HG0122110). An amino acid sequence alignment of all eight barley RIC proteins illustrated no general domain homologies outside the highly conserved CRIB motif (Fig. 1). Interestingly, the CRIB motif was more C-terminally located in RICs 153, 163 and 194, similar to RIC2 and RIC4 of *Arabidopsis thaliana* (Wu *et al.*, 2001), while the other RICs (157, 168, 170, 171 and 236) contained the CRIB motif closer to their N-terminal end. However, we didn’t identify additional conserved domains shared by all members of the barley RIC protein family.

**Figure 1:**
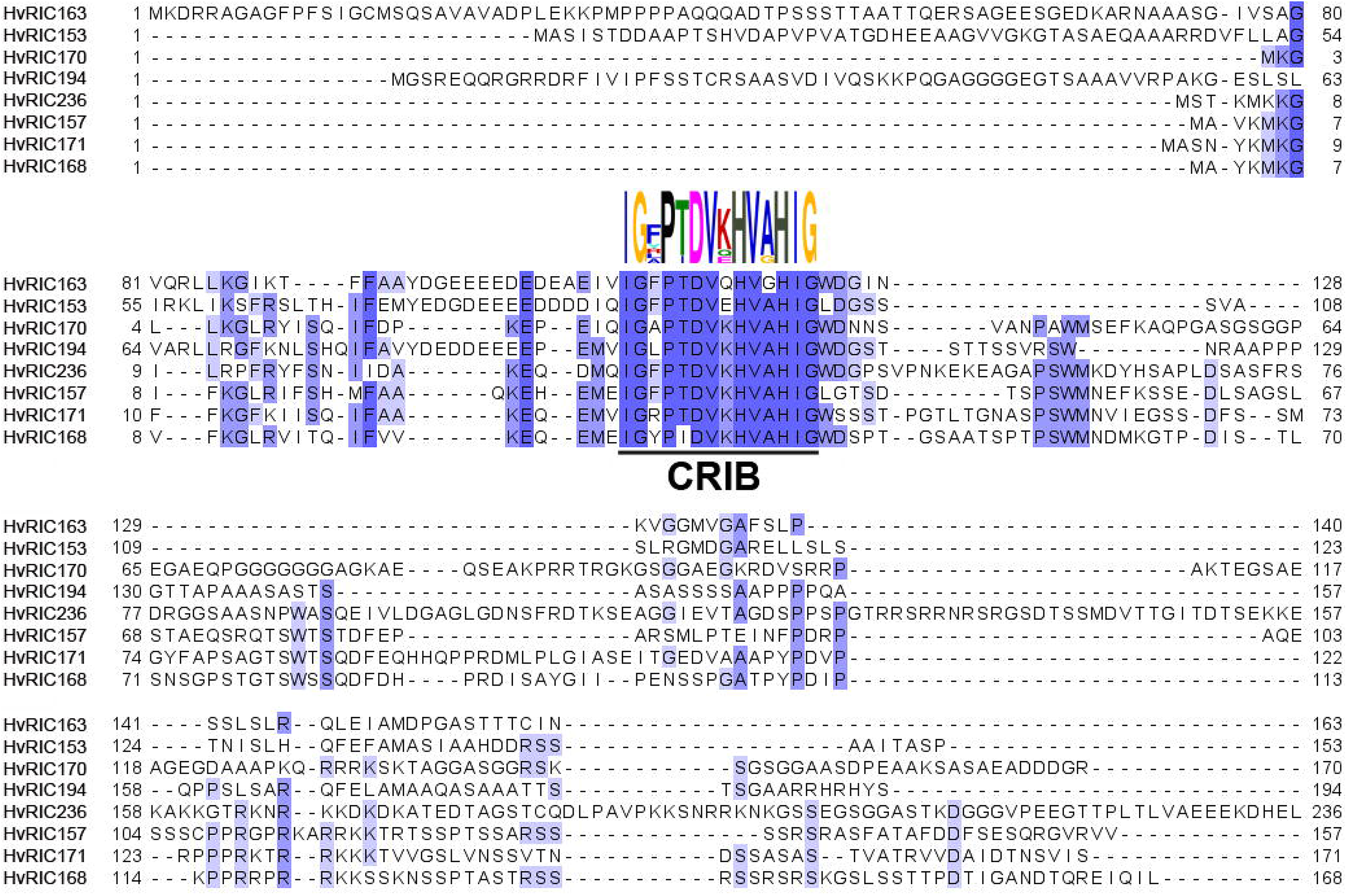
Barley RIC proteins alignment. Multiple alignment (https://www.ebi.ac.uk/Tools/msa/muscle, Edgar 2004; illustrated using Jalview 2.11.0 software) of predicted barley (*Hordeum vulgare*, *Hv*) proteins harboring CRIB domain (underlined). Intensity of blue coloured amino acids represents level of conservation (higher intensity = more conserved). CRIB domain consensus sequence of barley RIC proteins is indicated by coloured amino acid sequence above the alignment (http://meme-suite.org/tools/meme).

RACB-mediated susceptibility towards powdery mildew is determined exclusively in barley leaf epidermal cells. Hence, in order to unravel downstream signaling components of RACB, we focused on barley leaf-expressed RIC proteins. Using online available gene expression databases (https://webblast.ipk-gatersleben.de/barley_ibsc/), we identified five leaf-expressed *RIC* genes (*RIC153*, *RIC157*, *RIC163*, *RIC194* and previously published *RIC171* (Schultheiss *et al.*, 2008). Despite there is little sequence conservation between RIC proteins outside their CRIB domain, we compared primary sequences of barley, *Arabidopsis thaliana* and rice (*Oryza sativa*) RICs using an available online amino acid motif discovery software (http://meme-suite.org/tools/meme). This discovered three previously non-described amino acid motifs shared by indidual barley (RIC157, RIC168, RIC171), rice (Os02g06660.1, Os04g53580.1) and Arabidopsis RICs (RIC10, RIC11) (Suppl. Fig. S1) but not by all RIC proteins.

### RIC157 increases suspeptibility of barley towards Bgh in a RACB-dependent manner

To investigate a potential function of RIC157 during the barley-powdery mildew interaction, we analysed the penetration success of *Bgh* on barley epidermal cells during various conditions (Fig. 2). Single cell transient overexpression of RIC157 had a strong effect on the penetration success of *Bgh* into barley epidermal cells (Fig. 2A). The susceptibility to fungal cell entry increased by about 50% compared to control treatments, an outcome that is reminiscent of susceptibility levels observed after transient overexpression of constitutively activated CARACB(G15V) or RIC171 (Schultheiss *et al.*, 2003, Schultheiss *et al.*, 2008). We did not observe the opposite effect, meaning a decreased fungal penetration after RNA interference (RNAi)-mediated silencing of RIC157 compared to control levels (Fig. 2B). To check if this elevated susceptibility of barley epidermal cells after overexpression of RIC157 is dependent on RACB, we analysed the fungal penetration efficiency by simultaneous transient overexpression of RIC157 and silencing of endogenous RACB expression via RNAi (Fig. 2C). Interestingly, we did not observe an elevated fungal penetration rate, indicating that RIC157 increases barley epidermal cell susceptibility in a RACB-dependent manner. We confirmed the efficiency of both RNAi silencing constructs via co-expression of fluorescence tag-labelled targets and ratiometric fluorescence measurements (Suppl. Fig. S2).

**Figure 2:**
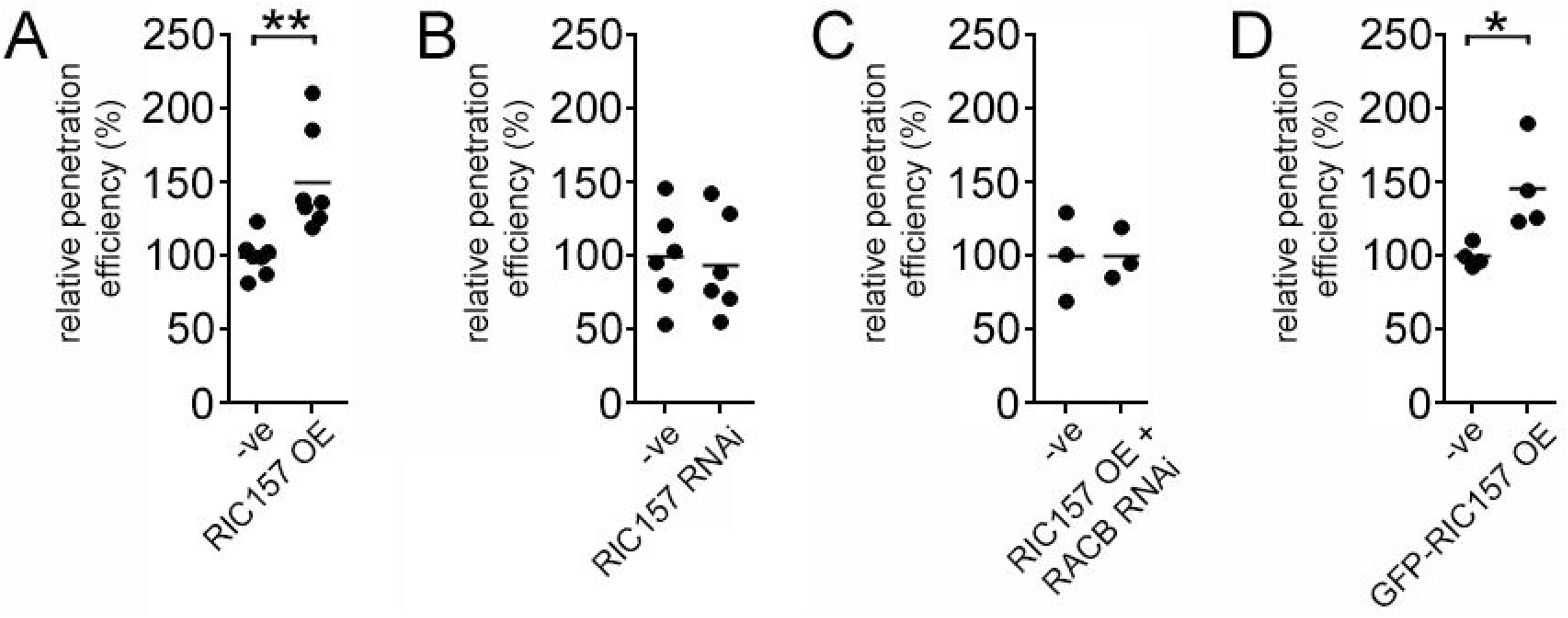
RIC157 increases susceptibility in a RACB-dependent manner. Epidermal cells of 7 day old primary barley leaves were transiently transformed by particle bombardment with either (A) an overexpression construct of RIC157, (B) a RNA interference construct of RIC157, (C) simultaneously with an overexpression construct of RIC157 and a RNA interference construct of RACB or (D) an overexpression construct of RIC157 N-terminally tagged with GFP. Empty overexpression or RNA interference plasmids were used as controls (−ve). The penetration efficiency of *Bgh* into transformed barley epidermal cells was analysed 48h after inoculation with fungal spores. Values are shown as mean of at least 3 independent biological replicates, relative to the mean of the control set as 100%. ** indicates significance P < 0.01, Student‘s t-test.

### RIC157 interacts directly with RACB in yeast and in planta

RACB can directly interact with CRIB motif-containing proteins RIC171 and MAGAP1 (Schultheiss *et al.*, 2008, Hoefle *et al.*, 2011). Therefore, it appeared likely that RIC157 can directly interact with RACB via its CRIB domain. We conducted different approaches to test for direct protein-protein interaction between RIC157 and RACB. In a targeted yeast-2-hybrid experiment, we showed that RIC157 directly interacts with RACB and with the constitutively activated CARACB(G15V) mutant, but not with the dominant-negative, GDP-bound DNRACB(T20N) mutant (Fig. 3A). Interestingly, we also observed a certain level of interaction between RIC157 and lower nucleotide affinity RACB mutant DNRACB(D121N). This particular mutation has been shown previously in RAS mutant D119N to behave either in a constitutively activated or a dominant-negative way, depending on the experimental setup (Cool *et al.*, 1999). The direct protein-protein interaction between RIC157 and RACB is dependent on the CRIB motif. RIC157 variants either lacking the CRIB motif completely or containing a CRIB motif that has been mutated at two highly conserved histidine residues (Burbelo *et al.*, 1995, Ash *et al.*, 2003), lose the ability to interact with either RACB form nearly entirely (Suppl. Fig. S3). To substantiate these results, we aimed to prove the fusion protein stability by immunoblotting (Suppl. Fig. S4). While all RACB variants were stably expressed and detectable, we were unable to confirm the stability of RIC157 variants in yeast in most of several independent experiments. Together with the fact that the yeast growth on selective medium was slow but dependent on the RIC157 construct, this suggests a high RIC157 turnover in yeast.

**Figure 3:**
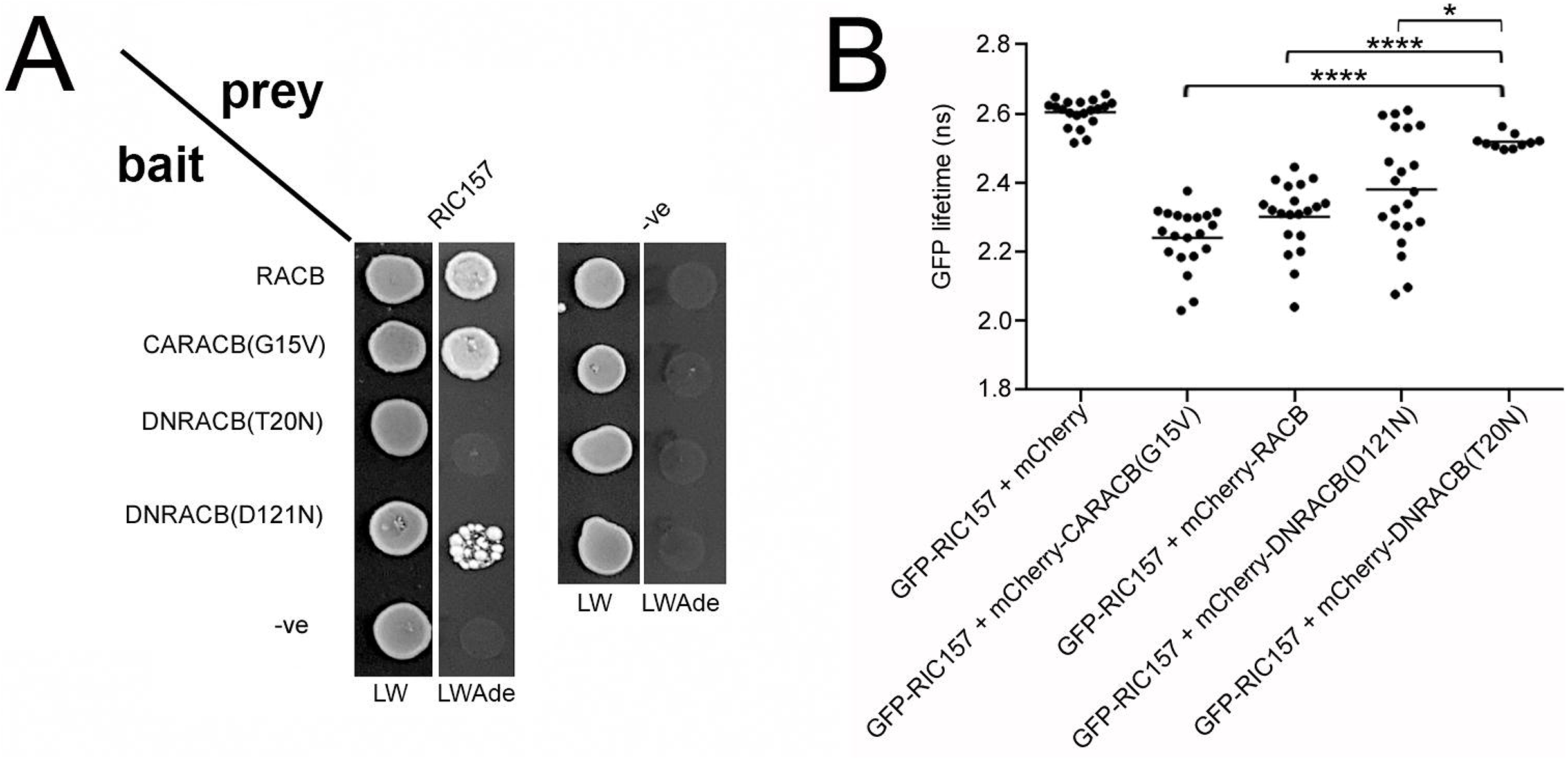
RIC157 interacts directly with RACB in yeast and in planta. A) Yeast-2-Hybrid indicates direct interaction between RIC157 and RACB in yeast. Yeast strain AH109 was transformed with indicated bait and prey fusion constructs. Overnight cultures of yeast transformants were dropped onto Complete Supplement Medium plates either lacking leucine and tryptophan (LW) or lacking leucine, tryptophan and adenin (LWAde) and incubated at 30°C. Growth on LWAde medium indicates interaction between bait and prey fusion proteins. –ve denotes empty prey and bait plasmid. Photos were taken 2 days (LW) and 7 days (LWAde), respectively, after dropping. B) Quantification of FLIM analysis confirms direct protein-protein interaction between RIC157 and RACB *in planta*. GFP lifetime in barley epidermal cells transiently co-expressing indicated constructs was investigated at the aequatorial plane 2d after transformation via particle bombardment. Graph shows result of 2 independent biological replicates. **** indicates significance P < 0.0001, Student‘s t-test.

In order to investigate the *in planta* interaction between RACB and RIC157, we performed Bimolecular Fluorescence Complementation (BiFC, synonym split-Yellow Fluorescent Protein (YFP)) experiments (Walter *et al.*, 2004). We therefore fused N-terminally N- and C-terminal YFP parts to RIC157 and RACB variants and transiently co-expressed complementary splitYFP fusion proteins in barley epidermal cells via particle bombardment. YFP fluorescence reconstitution was ratiometrically quantified against a co-expressed cytosolic mCherry fluorescence marker. As shown in Suppl. Fig. S5, YFP fluorescence was reconstituted to a significantly higher extent when split-YFP fusions of RIC157 were co-expressed with split-YFP fusions of RACB and CARACB(G15V), compared to co-expressions with DNRACB(T20N) and DNRACB(D121N). This suggests that RIC157 might preferantially interact with activated RACB *in planta*. We confirmed the stability of split-YFP fusion proteins via immunoblot analysis of total protein samples extracted from transformed barley mesophyll protoplasts (Suppl. Fig. S5B).

Since split-YFP experiments do not unequivocally reveal direct protein-protein interaction, we further analysed the interaction between RACB and RIC157 by FLIM-FRET and in particular the Green Fluorescent Protein (GFP) lifetime reduction (Fig. 3B). We fused GFP N-terminally to RIC157 and mCherry N-terminally to different RACB forms and transiently co-expressed respective combinations in barley epidermal cells via particle bombardment. Co-expression of GFP-RIC157 and free mCherry resulted in an average GFP lifetime of about 2.6 ns and is representative of a non-FRET negative control setup. GFP lifetime was slightly lower compared to the negative control samples in cells co-expressing GFP-RIC157 and mCherry-DNRACB(T20N). In contrast to that, the co-expression of GFP-RIC157 with mCherry-CARACB(G15V) led to a strong and highly significant reduction in GFP lifetime to approximately 2.2 ns. This GFP lifetime reduction clearly demonstrates a direct protein-protein interaction between RIC157 and activated RACB *in planta*. A very similar reduction in GFP lifetime we observed when GFP-RIC157 was co-expressed with mCherry-RACB suggesting an interaction of RIC157 with the wildtype form of RACB. The FLIM approach also indicated an interaction of RIC157 with the lower nucleotide affinity RACB mutant DNRACB(D121N). However, the spreading of the single data points was immense from no GFP lifetime reduction down to GFP lifetime reductions reminiscent of mCherry-CARACB(G15V)-expressing cells. Since the D121N mutation leads to a lower nucleotide affinity, this particular result we see here might have been provoked by different physiological cell conditions which in some cells stabilize the GDP-bound, inactive form and in others the GTP-bound, activated form of the DNRACB(D121N) mutant. In total, these different experimental approaches strongly suggest the direct protein-protein interaction between RIC157 and the activated form of RACB.

### Recruitment of RIC157 to the cell periphery is RACB dependent

To investigate the subcellular localisation of RIC157, we fused GFP N-terminally to RIC157 and transiently expressed this construct in barley epidermal cells via biolistic transformation. As shown in Figure 4A (upper row), RIC157 localises to the cytosol, but a lack of GFP fluorescence in the nucleoplasm indicated that GFP-RIC157 is excluded from the nucleus. However, we repeatedly observed a strong fluorescent signal around the nucleus suggesting a potential affinity of RIC157 to the nuclear envelope/endoplasmic reticulum membrane or proteins associated to it.

**Figure 4:**
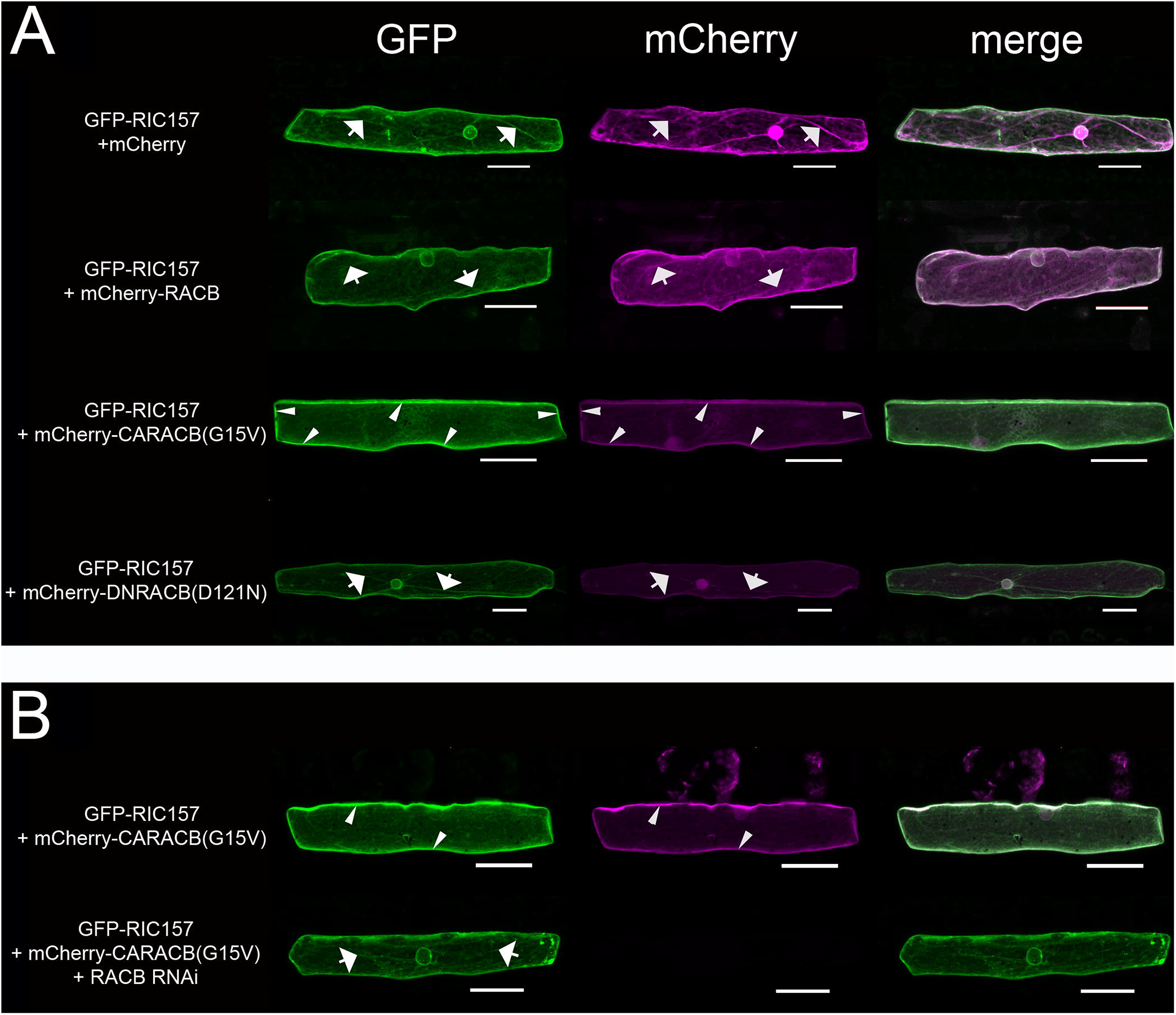
RIC157 localisation is affected by RACB-activation status. Confocal scanning microscopy of barley epidermal cells 1d after transformation via particle bombardment. A) GFP-RIC157 localises to the cytoplasm, but not the nucleus (upper row) and is recruited to the cell periphery exclusively by mCherry-CARACB(G15V), but not by mCherry-RACB or mCherry-DNRACB(D121N). B) Simultaneous RNA interference-mediated silencing of RACB attenuates RIC157 recruitment to the cell periphery. Arrows indicate cytoplasmic strands, arrow heads point towards GFP fluorescence accumulation at cell periphery. Microscopy pictures show maximum projections of at least 15 optical sections taken at 2μm increments. Bar = 50μm.

Since we found a direct protein-protein interaction between RIC157 and RACB, we checked the potential impact of RACB on the subcellular localisation of RIC157 by transiently co-expressing GFP-RIC157 with various untagged forms of RACB (Suppl. Fig. S6). The cytoplasmic localisation of RIC157 is not significantly affected in the presence of RACB or the lower nucleotide affinity dominant negative DNRACB(D121N) mutant. However, we observed a decrease in cytoplasmic and nuclear envelope-localised GFP fluorescence and an accumulation of GFP fluorescence at the cell periphery when GFP-RIC157 was co-expressed with the constitutively activated CARACB(G15V) form.

Because untagged proteins cannot be properly monitored, we extended our analysis in barley epidermal cells using mCherry-tagged RACB forms (Fig. 4A). Similarly, we detected GFP-RIC157 in the cytoplasm where it co-localised with mCherry fusions of RACB and DNRACB(D121N). In contrast to that, as with the untagged activated RACB, we detected a similarly strong re-localisation of GFP-RIC157 to the cell periphery in the presence of mCherry-tagged CARACB(G15V) that likewise accumulated at this site. This suggests that activated RACB, like other ROPs associating with the plasma membrane probably via its C-terminal prenylation and possible palmitoylation (Schultheiss *et al.*, 2003; Yalovsky 2015), recruits RIC157 to the cell periphery.

In order to check if this RIC157 recruitment to the cell periphery is indeed due to the co-expression with activated RACB, we simultaneously transformed barley epidermal cells with a RNAi construct to silence RACB (Fig. 4B). Without RACB silencing, co-expression of GFP-RIC157 and mCherry-CARACB(G15V) again lead to accumulation of both fusions proteins at the cell periphery. RNAi-mediated silencing of RACB on the other hand did not only diminish the mCherry fluorescence to almost non-detectable levels demonstrating RACB silencing took place, it also decreased GFP fluorescence at the cell periphery and increases GFP fluorescence in the cytoplasm. This result clearly supports that the observed recruitment of RIC157 to the cell periphery is mediated by activated RACB. To further confirm the recruitment of RIC157 by activated RACB from the cytoplasm to the plasma membrane, we took advantage of a mCherry-tagged plasma membrane marker, pm-rk (Nelson *et al.*, 2007), and analysed the potential co-localization of GFP-RIC157 with pm-rk in the absence and presence of co-overexpressed non-tagged constitutively activated RACB (CARACB(G15V), Suppl. Fig. S7). In the presence of activated RACB, fluorescence signals of GFP and mCherry overlapped to a higher extent compared to an experimental setup lacking overexpressed activated RACB. This unambiquously supports our previous findings that RIC157 is recruited by activated RACB to cell peripheral and plasma-membrane associated localisations.

Fluorescent proteins potentially have an impact on the functionality of the proteins to which they are fused due to conformational hindrances. In order to rule out that RIC157 fused to fluorescent proteins behaves differently than untagged RIC157, we analysed the penetration ability of *Bgh* in barley epidermal cells overexpressing GFP-tagged RIC157 (Fig. 2D). The powdery mildew fungus benefited from the presence of GFP-RIC157, similar to untagged RIC157, suggesting that GFP fusion to RIC157 did not prevent its ability to enhance susceptibility towards *Bgh* infection. Moreover, this supports that localisation of N-terminally tagged RIC157 proteins that we observe, represents the localization of a functional RIC157 protein.

### RIC157 and RACB co-localise and accumulate at penetration site

Co-localisation experiments in unchallenged barley epidermal cells demonstrated a recruitment of RIC157 to the cell periphery in the presence of activated RACB. In order to investigate if both proteins co-localise at a more specific subcellular site during the interaction with the powdery mildew fungus, we analysed the localisation of transiently co-expressed RIC157 and CARACB(G15V) in epidermal barley cells 18-24 hours after inoculation with *Bgh*. As shown in Fig. 5, fluorescent protein fusions of RIC157 and activated RACB accumulate close to the fungal penetration site, forming a cone outlining the neck of a developing haustorial initial. The RIC157/CARACB(G15V) co-localisation was much more defined than the fluorescence signal of simultaneously expressed cytoplasmic mCherry (Fig. 5B), indicating a specific membrane-associated co-localisation in epidermal cells that are successfully penetrated by the fungus.

**Figure 5:**
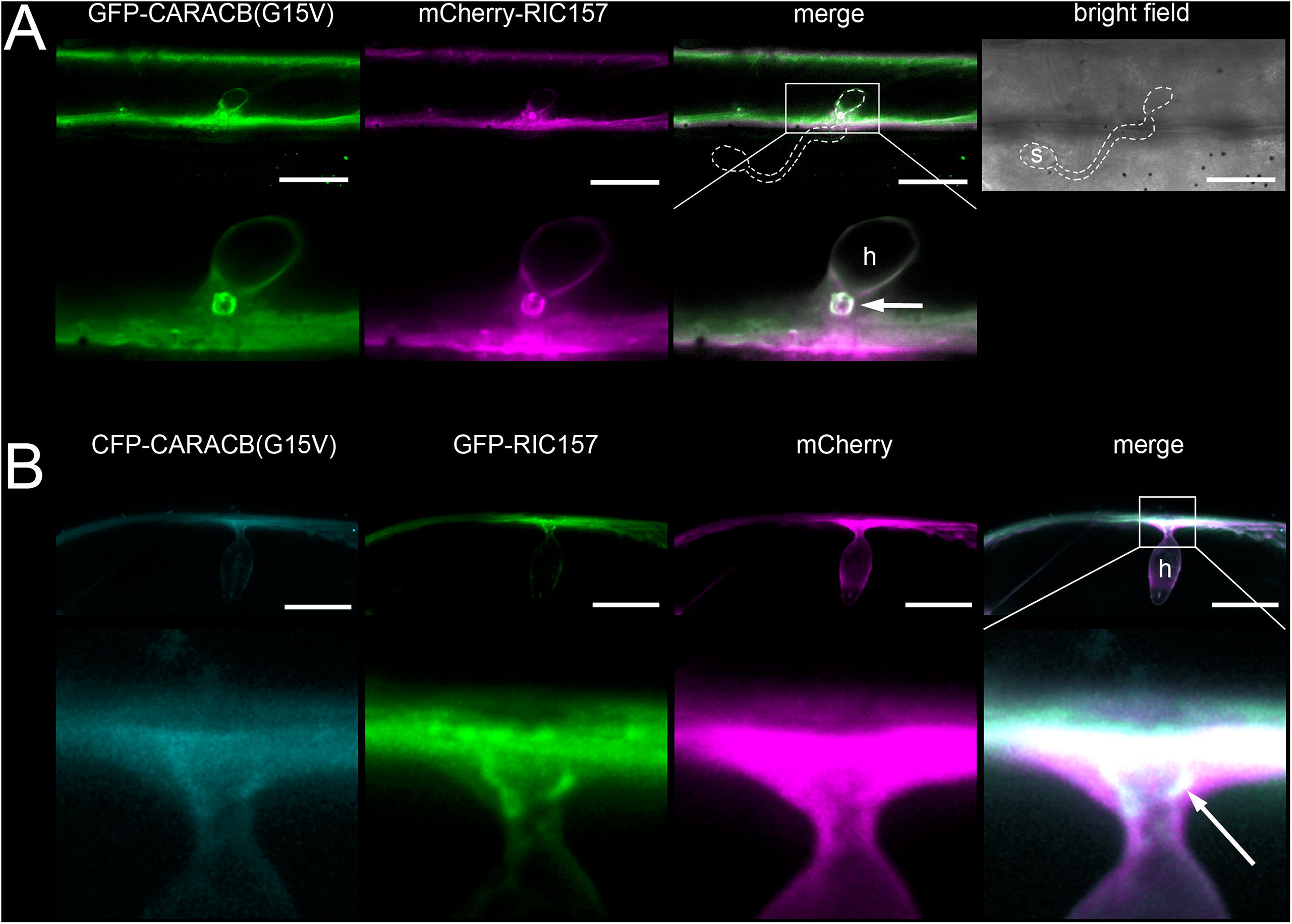
RIC157 is recruited to penetration site where it colocalises with activated RACB. Confocal laser scanning microscopy of epidermal cells 1d after transformation via particle bombardment and 18-24h after inoculation with *Bgh*. Cells in A) and B) show successful fungal penetration due to haustorium formation (h). A) Transient co-expression of GFP-CARACB(G15V) and mCherry-RIC157. Area in white square is enlarged in lower panel. S = spore; Bar = 30μm. B) Transient co-expression of CFP-CARACB(G15V) and GFP-RIC157. Cytosolic mCherry was expressed to distinguish RIC157 and CARACB(G15V) localisation from cytoplasm at penetration site. Bar = 20μm. Arrows indicate approximate position of the haustorial neck. Contrast of images was equally slightly enhanced. Arrows indicate haustorial neck close to fungal penetration site.

## Discussion

In the presence of *Bgh,* high activity of the ROP protein RACB appears to be disadvantageous for barley. To date, however, our knowledge about the exact RACB-regulated cellular processes, of which the fungus takes advantage, is still quite limited. Albeit we observed a role of RACB or RACB-interacting proteins in polar cell development and cytoskeleton organization (Opalski *et al.*, 2005; Hoefle *et al.*, 2011; Huesmann *et al.*, 2012; Scheler *et al.*, 2016; Nottensteiner *et al.*, 2018), a direct mechanistic link between RACB-mediated susceptibility and RACB-regulated cytoskeleton organisation or polar membrane trafficking is still missing. Our studies, however, open up the prospect of a RACB-regulated pathway via a ROP-specific scaffold protein that might be exploited in barley epidermal cells by *Bgh* to support susceptibility towards powdery mildew.

### RIC proteins as scaffolds in RACB downstream signalling

In a signaling cascade, scaffold proteins are as vital for mediating the molecular response as upstream signaling hubs or downstream executors. These scaffolds establish not just a hub-executor connection, by doing so they represent also the first branching point in the signalling cascade that eventually leads to specific effects without possessing any kind of enzymatic activity themselves (Zeke *et al.*, 2009, Good *et al.* 2011). Regarding signaling pathways, ROPs function as signaling hubs and have been shown to be involved in loads of cellular processes, and for some ROPs the regulatory role in different, sometimes even antagonistic signalling pathways has been described (Gu *et al.*, 2005, Nibau *et al.*, 2006, Feiguelman *et al.*, 2018). If ROPs do not interact directly with downstream executors, RIC proteins (and also ICRs/RIPs, Feiguelman *et al.*, 2018) are considered bridging units creating the scaffold for specific branches of ROP signaling (Schultheiss *et al.*, 2008, Craddock *et al.*, 2012, Zhou *et al.*, 2015, Hong *et al.*, 2016). Besides RIC171 and RIC157, we identified another six proteins of different sizes in barley that, by the above-mentioned definition, we consider RIC proteins. Potentially, there is a difference between monocots and dicots regarding the number of RIC proteins, which is slighthly higher in Arabidopsis (Wu *et al.*, 2001), suggesting a higher probability of either functional redundancies, diversification or antagonistic partners in dicots (Gu *et al.*, 2005). Redundancies between the barley leaf-expressed RIC proteins is, considering the low primary sequence conservation, hard to predict and not known yet. The partial similarities between leaf-expressed RIC157 and RIC171 (Suppl. Fig. 1) could, however, indicate functions in similar signaling pathways.

### RIC157 increases susceptibility towards powdery mildew

In this study, we concentrated on leaf-expressed RIC157 and show, that the transient RIC157 overexpression leads to a strong increase in barley epidermal cell susceptibility towards powdery mildew infection (Fig. 2A). Thus, the fungus benefits from a highly abundant RIC157. The RNA interference-mediated silencing of RIC157, however, did not lead to a higher resistance compared to control-treated cells (Fig. 2B). This could have different reasons. Although we have shown a significant reduction of ectopically expressed RIC157 protein levels in the presence of the RIC157 RNAi silencing construct (Suppl. Fig. S2), it must be noted that RNAi-based silencing is never 100% efficient. Remaining endogenous RIC157 transcript levels might be sufficient to allow for control level penetration efficiencies. Additionally, the protein turnover rate of RIC157 is unknown, meaning even with a high RNAi silencing efficiency it is still possible that RIC157 protein, expressed before transient transformation with the RNAi silencing construct, is present throughout the infection assay and sufficiently abundant to support penetration. Another very important point that needs to be stressed is the function of RACB or ROPs in general as signaling hubs (Nibau *et al.*, 2006). The signaling via RIC157 is likely not the only RACB-regulated path that leads to susceptibility. Shutting down the particular RACB-RIC157 route probably still leaves other RACB signaling branches functional, which are potentially involved to a certain extent in RACB-mediated susceptibility as well. An increased resistance towards fungal infection was, however, achieved once the signaling hub was removed or switched off by silencing RACB or by transient overexpression of its presumable antagonist MAGAP1 (Schultheiss *et al.*, 2002, Hoefle *et al.*, 2011). In accordance with this, RACB-dependency of the RIC157-promoted susceptibility (Fig. 2C) suggests that even abundant RIC157 still requires the presence of RACB to function as a susceptibility factor.

### RIC157 interacts with RACB and is recruited to the cell periphery

We have demonstrated that RIC157 can interact directly with RACB in yeast and *in planta* (Fig. 3, Suppl. Fig. 5). In barley epidermal cells, RIC157 showed interaction with activated RACB, but not with the dominant negative form. The observed interaction in yeast or via FLIM analysis *in planta* between RIC157 and the dominant-negative form DNRACB(D121N) could be explained by the different experimental setups and/or physiological cell conditions. G-proteins with this particular mutation have been observed previously to display either dominant-negative or constitutively activated properties (Cool *et al.*, 1999). DNRACB(D121N) has an intrinsic lower nucleotide affinity, however in yeast this form could be GTP bound and hence resemble the activated RACB.

The subcellular *in planta* CARACB(G15V)-RIC157 interaction site seen in BiFC and FLIM-FRET experiments appeared to be at the cell periphery. This seems conclusive, because RIC157 preferentially interacted with activated RACB. Activated ROPs are supposed to be associated with negatively charged phospholipids in plasmamembrane nanodomains, which is additionally promoted via posttranslational prenylation and S-acylation (Yalovsky 2015, Platre *et al.*, 2019). In *Arabidopsis thaliana* it has even been recently shown that particular phosphoinositides are recruited by filamentous pathogens to the plant-microbe interface (Qin *et al.*, 2020), suggesting a similar mechanism during powdery mildew infection of barley. Fluorescence-tagged RIC157 alone did not show any specific localisation in the absence of the fungus, although it seems to be excluded from the nucleus (Fig. 4). In the presence of activated RACB, however, RIC157 undergoes a relocalisation from the cytoplasm to the cell periphery/plasma membrane, where the interaction with the activated ROP takes place. It had previously been shown that expression of activated GFP-RACB alone leads to its preferential localisation at the cell periphery with a cytosolic background (Schultheiss *et al.*, 2003). We assume that a potential influence of endogenous levels of RIC157 or activated RACB on the localisation of their overexpressed interaction partner is probably negligible in our experimental setup. A similar recruitment to the cell periphery has been observed with other proteins that directly interact with activated RACB (Schultheiss *et al.*, 2008, Hoefle *et al.*, 2011, Huesmann *et al.*, 2012, McCollum *et al.*, 2019 Preprint), reinforcing the model that RACB activation is preceding the recruitment of interaction partners and that downstream RACB signaling is initiated at the plasma membrane. This is further supported since a RACB mutant version lacking the C-terminal CSIL motif for prenylation localizes to the cytoplasm and is inactive in promoting susceptibility (Schultheiss *et al.*, 2003).

With regard to RACB’s and RIC157’s capability to support fungal infection, the recruitment of RIC157 to the cell periphery becomes even more interesting. Our data suggest a recruitment of RIC157 to the fungal penetration site (Fig. 5). The cone-like structure surrounding the haustorial neck indicates a subcellular co-localisation with activated RACB. However, the co-localisation of RACB and fungal infection-supporting RACB-interactors is not exclusive to RIC157. RIC171 and RIPb, two proteins also considered scaffolds in RACB signalling, co-localise with RACB at the haustorial neck (Schultheiss *et al.*, 2008, Hückelhoven and Panstruga 2011, McCollum *et al.*, 2019 Preprint). Future experiments may show, whether subcellular co-concentration of RACB and RACB-interacting proteins indicate a specific lipid composition of the haustorial neck, which then recruits activated ROPs, or a membrane domain of high ROP activity due to local GEF activity, or perhaps indicates an exclusion of ROPs from further lateral diffusion into the EHM, which was suggested to be controlled at the haustorial neck (Koh *et al*., 2005).

### RIC157 and susceptibility

RIC157 transiently overexpressed in barley epidermal cells localises to the cytoplasm and enhances fungal penetration efficiency in a RACB-dependent manner. Activated RACB, however, associates with the plasma membrane where it likely recruits RIC157 for downstream signaling. Thus, transiently overexpressed and endogenous RIC157 might promote RACB-mediated susceptibility upon recruitment to the cell periphery by endogenously present activated RACB. Only a small fraction of overexpressed RIC157 is possibly recruited to the cell periphery without concomitant co-overexpression of activated RACB. This means that such a minute fluorescence localisation change might be probably undetectable in our experimental setup, but it needs to be emphasized that we do not assume RIC157 promoting susceptibility to powdery mildew from a solely cytoplasmic site. Indeed, in front of a cytoplasmic background, RIC157 is also visible at the cell periphery without co-expression of CARACB(G15V), possibly reflecting partial recruitment by endogenous ROPs (Fig. 4A).

The domain and sequence similarity between barley RIC157 and *Arabidopsis thaliana* RIC10 and RIC11 (Suppl. Fig. 1) does not necessarily indicate similar functions of both proteins. Beside AtRIC10 and AtRIC11, for which nothing is known to date about their biological function, RIC157 also shares limited amino acid motif similarity with AtRIC1, AtRIC3 and AtRIC7 (Suppl. Fig. 8), for which an involvement in cytoskeleton organization has previously been demonstrated. From these three Arabidopsis RIC proteins, AtRIC1 appears to be an interesting candidate from which a RIC157 function could be deduced. AtRIC1 has been shown to interact with ROP6 to activate the p60 subunit of Katanin, a microtubule-severing enzyme (Lin *et al.*, 2013), as opposed to the interaction with AtROP2 that negatively regulates the action of AtRIC1 on microtubules (Fu *et al.*, 2005). The microtubule-severing activity of Katanin might even be regulated by AtROP2, AtROP4 and AtROP6 (Ren *et al.*, 2017) via AtRIC1. Whether barley RIC157 also fulfils such a regulatory role in organising microtubules still remains to be seen. Regarding the subcellular localisation there are, however, clear differences: Barley RIC157 localises to the cytoplasm, AtRIC1 associates with microtubules (Fu *et al.*, 2005). However, when microtubule arrays in penetrated and attacked but non-penetrated barley epidermal cells were compared, penetration success was strongly associated with parallel non-polarized microtubule arrays in the cell cortex and a diffuse or depleted microtubule structure at the haustorial neck (Hoefle *et al.*, 2011). AtRIC3 has been shown to be involved in the pollen tube growth process, where its function leads to actin disassembly in a ROP1-dependent manner upon calcium influx into the cytoplasm (Gu *et al.*, 2005, Lee *et al.*, 2008). Likewise, AtRIC7 was recently reported to influence vesicle trafficking in stomata resulting in the suppression of an elevated stomatal opening after ROP2-dependent inhibition of the exocyst complex *via* Exo70B1 (Hong *et al.*, 2016). Both F-actin organization and exocyst function are important in penetration resistance to *Bgh* (Opalski *et al.*, 2005, Miklis *et al.*, 2007, Ostertag *et al.*, 2012). Therefore, the discovery of RIC157 as a RACB-dependent susceptibility factor may pave the way to a better understanding of ROP-steered processes that are pivotal for fungal invasion into barley epidermal cells. The challenge will be to find the downstream factors that RIC157 activates and to understand how RIC157 interacts with RACB in the presence of several other RACB interactors. We assume that several diverse RICs and ICR/RIP proteins could form a cooperative network for orchestrating F-actin, microtubule and membrane organization at the site of fungal entry.

## Experimental procedures

### Plant and fungal growth conditions

Wildtype barley (*Hordeum vulgare*, cultivar „Golden Promise“) was cultivated in long day conditions (16 hours day light, 8 hours darkness) at a temperature of 18°C with a relative humidity of 65% and a light intensity of 150μmol s^−1^ m^−2^.

The biotrophic powdery mildew fungus *Blumeria graminis* f.sp. *hordei* A6 was used in all experiments. It was cultivated and propagated on barley „Golden Promise” under the same condition described above.

### Cloning of constructs

Via a Two-Step Gateway cloning approach, in a first PCR *RIC157* (HORVU.MOREX.r2.6HG0469110) was amplified from a barley cDNA pool prepared from leafs and epidermal peels using gene-specific primers RIC157_GW_for and RIC157_GW_rev+STOP (Suppl. Table 1) creating an incomplete Gateway attachment site overhang. A second PCR using primers attB1 and attB2 completed the attachment sites. To create a Gateway entry clone of *RIC157*, the amplified product was recombined into pDONR223 (Invitrogen) via BP-reaction using Gateway BP Clonase™ ∥ according to manufacturer’s instruction (Thermo Fisher Scientific). To clone *RIC157ΔCRIB*, both fragments upstream and downstream of CRIB motif were amplified seperately using primers RIC157_GW_for and RIC157delCRIB_rev for PCR1, RIC157delCRIB_for and RIC157_GW_rev+STOP for PCR2, creating overlapping overhangs. In PCR3 both fragments together with primers RIC157_GW_for and RIC157_GW_rev+STOP completed *RIC157ΔCRIB* with incomplete Gateway attachment sites overhangs. Completetion of attachment sites and creating an entry clone in pDONR223 was done as described above. To clone *RIC157-H37Y-H40Y*, a site-directed mutagenesis using primers CRIB157H37&40Y_for and CRIB157H37&40Y_rev was performed according to QuikChange® Site-Directed Mutaganesis Protocol (Stratagene). For cloning entry constructs of *RACB* variants, primers RACB_GW_for and RACB_GW_rev were used to amplify *RACB* from previously described constructs (Schultheiss *et al.*, 2003) and cloned into pDONR223 via BP as described abobe. To clone *DNRACB(D121N)*, a site-directed mutagenesis using primers RACB_D121N_fw and RACB_D121N_rv was performed according to QuikChange® Site-Directed Mutaganesis Protocol (Stratagene).

For RNA interference (RNAi) silencing of *RIC157* in barley, we PCR-amplified two RIC157 fragments, a 97bp fragment with primers RIC157_RNAi_NotI_for and RIC157_RNAi_EcoRI_rev containing NotI and EcoRI restriction sites, and a 324bp fragment with primers RIC157_RNAi_EcoRI_for and RIC157_RNAi_XbaI_rev containing EcoRI and XbaI restriction sites. After restriction digest of all sites, ligating into pIPKTA38 via NotI and XbaI sites we created a *RIC157*_RNAi entry construct lacking the CRIB motif nucleotide sequence to prevent off-target silencing of other CRIB-domain containing RNAs. The *RIC157*_RNAi sequence was then cloned via LR reaction using Gateway LR Clonase™ II according to manufacturer’s instruction (Thermo Fisher Scientific) into RNAi expression plasmid pIPKTA30N to create a double-strand RNAi expression construct (Douchkov *et al.*, 2005).

For Yeast-2-Hybrid expression clones, entry clones of *RIC157* and *RACB* variants were introduced into prey plasmid pGADT7-GW and pGBKT7-GW via LR reaction using Gateway LR Clonase™ II according to manufacturer’s instruction (Thermo Fisher Scientific). pGADT7-GW and pGBKT7-GW have been modified from pGADT7 and pGBKT7 (Clontech) into a Gateway-compatible form using Gateway™ Vector Conversion System (Thermo Fisher Scientific). To create *RACB* variants lacking C-terminal prenylation sequence, a premature STOP-Codon was introduced by site-directed mutagenesis as described above using primers delCSIL_for and delCSIL_rev.

In order to clone BiFC constructs, we PCR amplified a *RIC157* full-length fragment with primers RIC157_BamHI_for and RIC157_KpnI_rev containing BamHI and KpnI restriction sites. After restriction digest, we ligated this construct into pUC-SPYNE(R)173 (Waadt *et al.*, 2008).

For localisation and overexpression studies in barley, *RIC157* and *RACB* variants in pDONR223 were used as entry constructs to clone them into various pGY1-based CaMV35S promoter-driven expression vectors (Schweizer *et al.*, 1999) via LR reaction as described above. Empty pGY1 (encoding for no tag) was rendered Gateway-compatible via Gateway™ Vector Conversion System (Thermo Fisher Scientific). To create expression vectors for proteins C- or N-terminally tagged by GFP or mCherry, Gateway Reading Frame Cassettes for C- and N-terminal fusions, respectively, were integrated into a pGY1-plasmid backbone upon XbaI digestion and combined at 5‘ or 3‘ with sequences of monomeric GFP or mCherry. Cloning procedure was performed using In-Fusion HD cloning kit (Takara Bio USA). Constructs for GFP and mCherry upstream or downstream of the Gateway cassette were amplified using primers GW_RfA_mCherry-F, GW_RfA_meGFP-F, GW_RfA_Xba-R, GW_Xba_RfB-F, GW_RfB-R, meGFP-STP-F, mCherry-STP-F, XFP-noSTP_Xba-F, XFP-noSTP-R, meGFP-noSTP-R, mCherry-STP_Xba-R and meGFP-STP_Xba-R.

The RACB RNAi construct, RACB BiFC constructs have been described previously (Schultheiss *et al.*, 2003, Schultheiss *et al.*, 2008, Schnepf *et al.*, 2018, McCollum *et al.*, 2019 Preprint).

### Barley epidermal cell transformation and penetration efficiency assessment

For transient overexpression in barley, primary leaf epidermal cells of 7d old plants were transformed using biolistic bombardment with 1μm gold particles that were coated with 2μg of each test plasmid and additionally with 1μg of a cytosolic transformation marker. After mixing the gold particles with plasmid combinations, CaCl_2_ (0.5M final concentration) and 3.5μl of 2mg/ml Protamine (Sigma) were added to each sample. The gold particle solution was incubated at room temperature for 30min, washed twice with 500μl Ethanol (first 70%, then 100%) and eventually dissolved in 6μl 100% ethanol per biolistic transformation. After shooting, leaves were incubated at 18°C.

For localisation and BiFC experiments, leaves were analysed 2 days after transformation. For FRET-FLIM analysis of RACB-RIC157 interaction, barley primary leaves of 7d old plants were transiently transformed. Therefore, 2ug of mCherry-RACB and 1ug meGFP-RIC157 containing plasmids were coated on gold particles for biolistic transformation of single barley epidermal cells.

For inoculation with *Bgh*, fungal spores were manually blown in a closed infection device over transformed leaves either 6 hours after transformation (for microscopic analyses 16 hours after inoculation) or 1 day after transformation (to check penetration efficiency 48 hours after inoculation).

To analyse penetration efficiency, a transient assay system based on a cytosolic GUS marker was used as decribed previously (Schweizer *et al.*, 1999). The reporter gene construct pUbiGUSPlus was a gift from Claudia Vickers (Addgene plasmid # 64402; http://n2t.net/addgene:64402; RRID:Addgene_64402, Vickers *et al.*, 2003). Additionally to overexpression or RNAi silencing constructs, each barley leaf was co-transformed with pUbiGUSPlus. 48 hours after *Bgh* inoculation, leaves were submerged in GUS staining solution (0.1M Na_2_HPO_4_/NaH_2_PO_4_ pH 7.0, 0.01 EDTA, 0.005M Potassium hexacyanoferrat (II), 0.005M Potassium hexacyanoferrat (III), 0.1% (v/v) Triton X-100, 20% (v/v) Methanol, 0.5mg/mL 1.5-bromo-4-chloro-3-indoxyl-β-D-glucuronic acid). For the solution to enter the leaf interior, a vacuum was applied. The leaves were incubated at 37°C over night in GUS staining solution and subsequently for at least 24 hours in 70% Ethanol. Fungal structures were stained with ink-acetate solution (10% ink, 25% acetic acid). Transformed cells were identified after GUS staining with light microscopy. An established haustorium was considered a successful penetration and for each sample at least 50 interactions were analysed. Barley epidermal cells transformed with the empty expression plasmid were used as negative control.

### Barley protoplast preparation and transformation

To prepare protoplasts from barley mesophyll cells, the lower epidermis of primary leaves from 7 day-old barley plants was peeled and the leaves were incubated 3 to 4 hours at room temperature in the darkness while floating with the open mesophyll facing downwards on an enzymatic digestion solution: 0.48M mannitol, 0.3% (w/v) Gamborg B5, 10mM MES pH 5.7, 10mM CaCl_2_, 0.5% (w/v) Cellulase R10, 0.5% (w/v) Driselase, 0.5% Macerozyme R10. After enzymatic treatment, an equal amount of W5 solution was added: 125mM CaCl_2_, 154mM NaCl, 5mM KCl, 2mM MES pH 5.7. Upon filtering through a 40μm nylon mesh, the protoplasts were pelleted 5min at 200g and carefully resuspended in 10ml W5 solution. After another centrifugation step, the protoplast concentration was adjusted to 2 x 10^6^ cells per mL in MMG solution: 0.4M mannitol, 15mM CaCl_2_, 2mM MES pH 5.7. For each transformation sample, 1mL protoplast solution was mixed with 50μg of each plasmid and 1.1mL PEG solution (40% (w/v) PEG4000, 0.1M mannitol, 0.2M CaCl_2_) and incubated 20min at room temperture in the darkness. Afterwards, 4.4mL of W5 solution was added to each transformation and gently mixed. After another pelleting at 200g, the protoplasts were resuspended in 1mL W1 solution (0.5M mannitol, 20mM KCL, 4mM MES pH 5.7) and incubated in the darkness at room temperature for at least 16 hours.

### Yeast-2-Hybrid

Yeast strain AH109 was transformed with bait (pGBKT7) and prey (pGADT7) constructs by following the small scale yeast transformation protocol from Yeastmaker™ Yeast Transformation System 2 (Clontech). Upon transformation, yeast cells were plated on Complete Supplement Medium (CSM) plates lacking leucine and tryptophan (LW) and incubated for 3 days at 30°C. A single colony was taken to inoculate 5mL of LW-dropout liquid medium that was incubated with shaking over night at 30°C. The next day, 2mL of culture was pelleted for immunoblot analyses. 7.5μL of undiluted overnight culture (and additionally a 1:10, 1:100 and 1:1000 for control purposes) were dropped on CSM plates lacking leucine and tryptophan, and also on CSM plates lacking leucine, tryptophan and adenine. Plates were incubated for at least 3 days at 30°C. Growth on CSM-LW plates confirmed the successful transformation of both bait and prey plasmids, while growth on CSM-LWAde plates indicated activation of reporter genes. As control for a positive and direct protein-protein interaction we routinely used murine p53 and the SV40 large T-antigen (Li and Fields 1993).

### Immunoblot analysis

For total protein extraction from yeast, we followed the protocol described in Kushnirov (2000) using 2mL of over night culture of yeast transformants grown in CSM-LW liquid medium. The extraction of total protein from barley mesophyll protoplasts was performed by pelleting transformed protoplasts 5min at 200g. The pellet was resuspended thoroughly in 50μL of 2x SDS loading buffer by vortexing. A complete protein denaturation was achieved by boiling protoplast samples 10min at 95°C. After shortly spinning down the samples, the stability of fusion proteins in yeast and in planta was assessed via Sodium dodecylsulfate-polyacrylamide gel electrophoresis (SDS-PAGE) and immunoblotting on PVDF membranes. Antibodies used for detecting protein bands on PVDF membranes came from SantaCruz Biotechnology (https://www.scbt.com/scbt/cart/cart.jsp): anti-GFP(B-2), anti-cMyc(9E10), anti-HA(F-7) and horseradish-peroxidase conjugated anti-mouse. Presence of antibodies on membrane was visually detected by using SuperSignal West Femto Chemiluminescence substrate (ThermoFisher Scientific). Equal protein loading and blotting success was confirmed via Ponceau S-staining of the PVDF membrane.

### Microscopy

Localisation and BiFC experiments were analysed on a Leica TCS SP5 confocal laser scanning microscope. The excitation laser wavelengths were 458nm for CFP, 488nm for GFP, 514nm for YFP and 561nm for RFP and mCherry, respectively. The fluorescence emission was collected from 462 to 484nm for CFP, from 500-550nm for GFP, from 515 to 550nm for YFP and from 569 to 610nm for RFP and mCherry. Barley epidermal cells were imaged via sequential scanning as z-stacks in 2μm increments. Maximum projections of each z-stack were exported as Tiff files from the Leica LAS AF software (version 3.3.0).

Localisation experiments of fluorescent protein fusions and BiFC analysis in barley epidermal cells were conducted from 24 hours until 48 hours after biolistic transformation. Regarding raciometric BiFC quantification using Leica LAS AF software (version 3.3.0), the fluorescence intensity was evaluated over the whole cell area and the ratio between YFP and cytosolic mCherry fluorescence signal was calculated. For each BiFC combination, the fluorescence of at least 20 cells was measured.

For FRET-FLIM analysis of RACB-RIC157 interaction, the expression of the fluorophore-fusion proteins was analysed 1 day after transformation using an Olympus FluoView™3000 inverse laser scanning confocal microscope with an UPLSAPO 60XW 60x/NA 1.2/WD 0.28 water immersion objective (Olympus, Hamburg, Germany). Fluorescence of GFP was collected between 500-540nm and mCherry emission was imaged between 580-620nm upon excitation with 488 and 561nm argon laser lines, respectively. For FRET-FLIM measurements the PicoQuant advanced FCS/FLIM-FRET/rapidFLIM upgrade kit (PicoQuant, Berlin, Germany) was used, comprising a 485nm pulsed laser line for GFP excitation (pulse rate 40 mHz, laser driver: PDL 828 SEPIA II, laser: LDH-D-C-485), a Hybrid Photomultiplier Detector Assembly 40 to detect GFP fluorescence and a TimeHarp 260 PICO Time-Correlated Single Photon Counting module (resolution 25 ps) to measure photon life times. GFP fluorescence was imaged at the aequatorial plane of epidermis cells to capture GFP fluorescence at the cell periphery and possibly plasma membrane. For each interaction at least 10 cells were analysed in two replicates and a minimum of 1000 photons of the brightest pixel were recorded. Decay data within a region of interest were fitted using an n-exponential reconvolution fit with model parameters n=3 and measured instrument response function.

## Acknowledgements

This work was supported by research grants from the German research foundatation (HU886/8 and SFB924) to RH. We are grateful to the TUM plant technology center for support in propagation of barley plants.

## Supplemental Data

**Suppl. Fig. S1: Barley RIC157 shows limited sequence similarity to RIC1 from *Arabidopsis thaliana***

**Suppl. Fig. S2: RNA interference silencing efficiency**

**Suppl. Fig. S3: CRIB deletion and CRIB mutation in RIC157 prevents interaction with RACB in yeast**

**Suppl. Fig. S4: Protein stability in yeast**

**Suppl. Figure S5: Raciometric BiFC analyses and BiFC fusion protein stability**

**Suppl. Fig. S6: RIC157 is recruited to the cell periphery by activated RACB**

**Suppl. Fig. S7: RIC157, recruited to the cell periphery by activated RACB, colocalises with plasma membrane marker.**

**Suppl. Fig. S8: Barley RIC157 shows limited sequence similarity to RIC proteins from *Arabidopsis thaliana*.**

**Suppl. Table 1: Primers used in this study**

---

**Suppl. Fig. S1: Barley RIC157 shows limited sequence similarity to barley RIC168 and RIC171, but also to RIC proteins from *Arabidopsis thaliana* and *Oryza sativa***. Primary sequences of RIC proteins (schematically shown in top part of the figure) were analysed for domain homologies using MEME online software (http://meme-suite.org/tools/meme). Consensus sequence of the four most similar motifs are shown below. Underlined sequence denotes CRIB domain. Hv = *Hordeum vulgare*, At = *Arabidopsis thaliana*, Os = *Oryza sativa*.

**Suppl. Fig. S2: RNA interference-mediated silencing efficiency.** Epidermal cells of 7d old barley primary leaves were transiently transformed via particle bombardment with overexpression constructs of GFP fusions of RIC157 (A) and RACB (B) alone or together with RNAi silencing constructs. A construct to express cytosolic mCherry was simultaneously co-delivered for transformation efficiency and fluorescence quantification purposes. Microscopy images are maximum projections of at least 15 optical sections taken at 2μm increments. Bars = 50μm. Each graph shows the mean of GFP fluorescence as percentage of mCherry fluorescence per transformed cell (whole cell area was taken as region of interest for measuring fluorescence intensity). Dots represent single measured cells. **** indicates significance P < 0.0001

**Suppl. Fig. S3: CRIB deletion and CRIB mutation in RIC157 prevents interaction with RACB in yeast.** Yeast strain AH109 was transformed with indicated bait and prey fusion constructs. Overnight cultures of yeast transformants were dropped onto Complete Supplement Medium plates either lacking leucine and tryptophan (LW) or lacking leucine, tryptophan and adenin (LWAde) and incubated at 30°C. Growth on LWAde medium indicates interaction between bait and prey fusion proteins. –ve denotes empty prey and bait plasmid. Photos were taken 2 days (LW) and 7 days (LWAde), respectively, after dropping.

**Suppl. Fig. S4: Protein stability in yeast.** Immunoblots probed with α-cMyc to detect RACB variants fused to GAL4 binding domain (BD; bait) encoded on pGBKT7 and α-HA to detect RIC157 variants fused to GAL4 activation domain (AD; prey) encoded on pGADT7 after transformation of yeast strain AH109. Total yeast protein was extracted as described in Experimental Procedures. Molecular weight is indicated in kiloDalton (kDA). Ponceau S staining shows protein loading. Stable expression of RIC157 variants fused to GAL4 activation domain could not be detected in immunoblots (arrow indicates expected bandsize).

**Suppl. Figure S5: Raciometric BiFC analyses and BiFC fusion protein stability.** A) Barley epidermal cells were transiently transformed with indicated constructs encoding for BiFC fusion proteins. Cytosolic mCherry expression was used in each sample as transformation marker and to quantify reconstituted YFP fluorescence ratiometrically. The whole cell area was taken as region of interest. Graph shows the mean of the YFP/mCherry fluorescence ratio, taken from at least 20 cells (shown as dots. **** indicates significance P < 0.0001, ** indicates significance P < 0.01 Student’s t-test. B) Barley mesophyll protoplasts were prepared and transformed with BiFC constructs as described in Experimental procedures. To confirm stable *in planta* expression, immunoblots were probed with α-HA to detect YFPc-RACB fusion proteins and α-cMyc to detect YFPn-RIC157 fusion proteins. Ponceau S staining shows equal protein loading. C) BiFC shows close proximity between RIC157 and RACB *in planta*. Barley epidermal cells transiently co-expressing split-YFP fusion protein combinations and mCherry as cytosolic transformation marker after particle bombardment transformation. 2d after transformation, YFP fluorescence reconstitution was analysed via Confocal laser scanning microscopy. Shown are maximum projections of at least 15 optical sections taken at 2μm increments. Bars = 50μm.

**Suppl. Fig. S6: RIC157 is recruited to the cell periphery by activated RACB.** Confocal laser scanning microscopy of barley epidermal cells 1d after transient transformation via particle bombardment. GFP-RIC157 localises to the cytoplasm, but not the nucleus and is recruited to the cell periphery exclusively by non-tagged CARACB(G15V), but not DNRACB(D121N). A construct for cytosolic mCherry expression was simultaneously used to check transformation efficiency. Arrows indicate cytoplasmic strands, arrow heads point towards GFP fluorescence accumulation at cell periphery. Microscopy pictures show maximum projections of at least 15 optical sections taken at 2μm increments. Bar = 50μm.

**Suppl. Fig. S7: RIC157, recruited to the cell periphery by activated RACB, colocalises with plasma membrane marker.** Confocal laser scanning microscopy of barley epidermal cells 1d after transient transformation via particle bombardment to express GFP-RIC157, red-fluorescent plasma membrane marker pm-rk (Nelson *et al.*, 2007) and non-tagged CARACB(G15V). Microscopy pictures show A) maximum projections of at least 15 optical sections taken at 2μm increments to visualize global recruitment of GFP-RIC157 to the cell periphery in the prescence of activated RACB or B) and C) a single optical section to confirm specific colocalisation of GFP-RIC157 with plasma membrane marker pm-rk. B) and C) show overlay analysis of both fluorescences in the absence or presence of overexpressed CARACB(G15V), either via magnification (inset in upper panel) or via graphical visualization of fluorescence signal intensities (lower panel) measured over a region of interest (white line in middle panel). Bar = 50μm.

**Suppl. Fig. S8: Barley RIC157 shows limited sequence similarity to RIC proteins from *Arabidopsis thaliana***. Primary sequences of RIC proteins (schematically shown in top part of the figure) were analysed for domain homologies using MEME online software (http://meme-suite.org/tools/meme). Consensus sequence of the four most similar motifs are shown below. Underlined sequence denotes CRIB domain. Hv = *Hordeum vulgare*, At = *Arabidopsis thaliana*.

**Suppl. Table 1:**
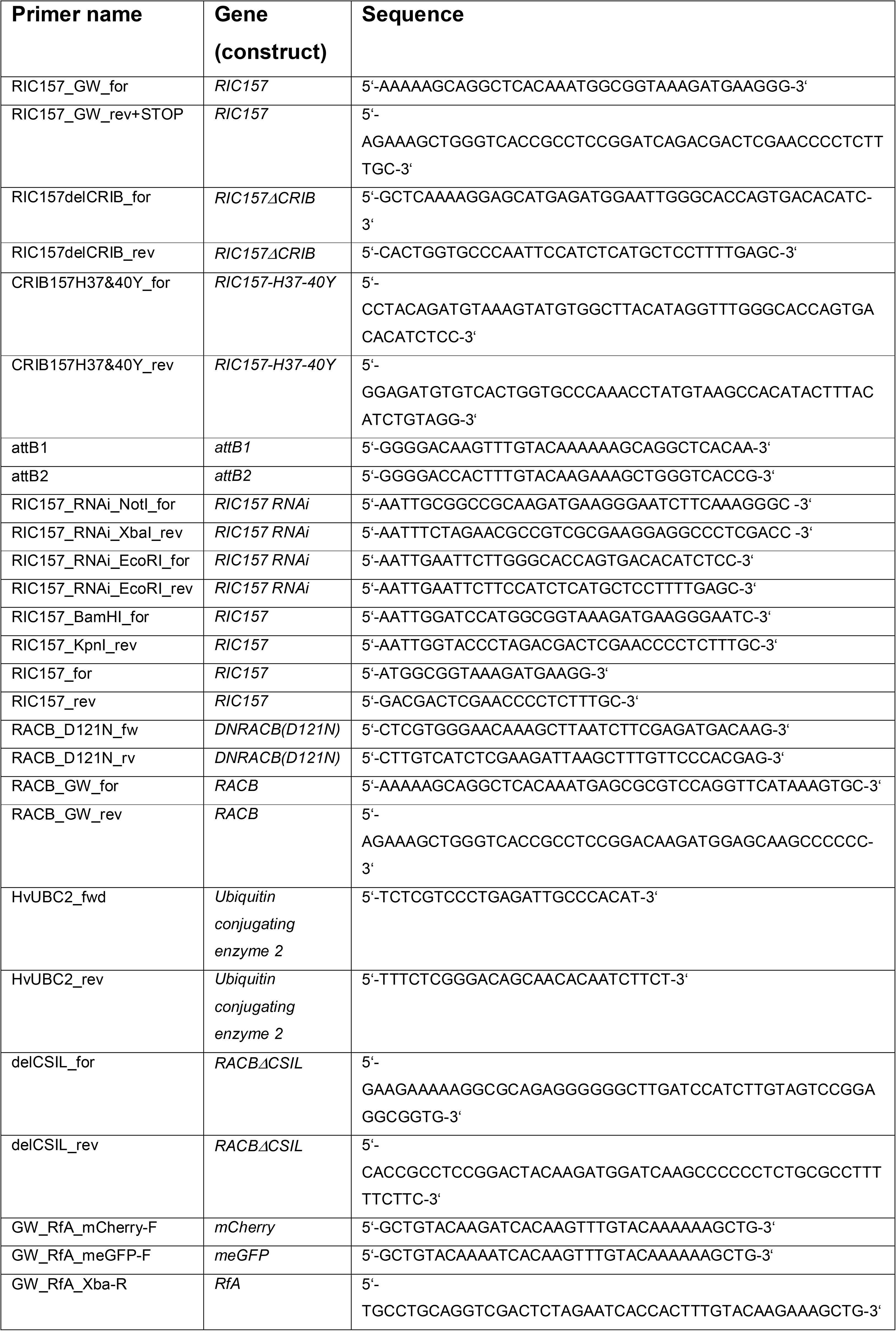
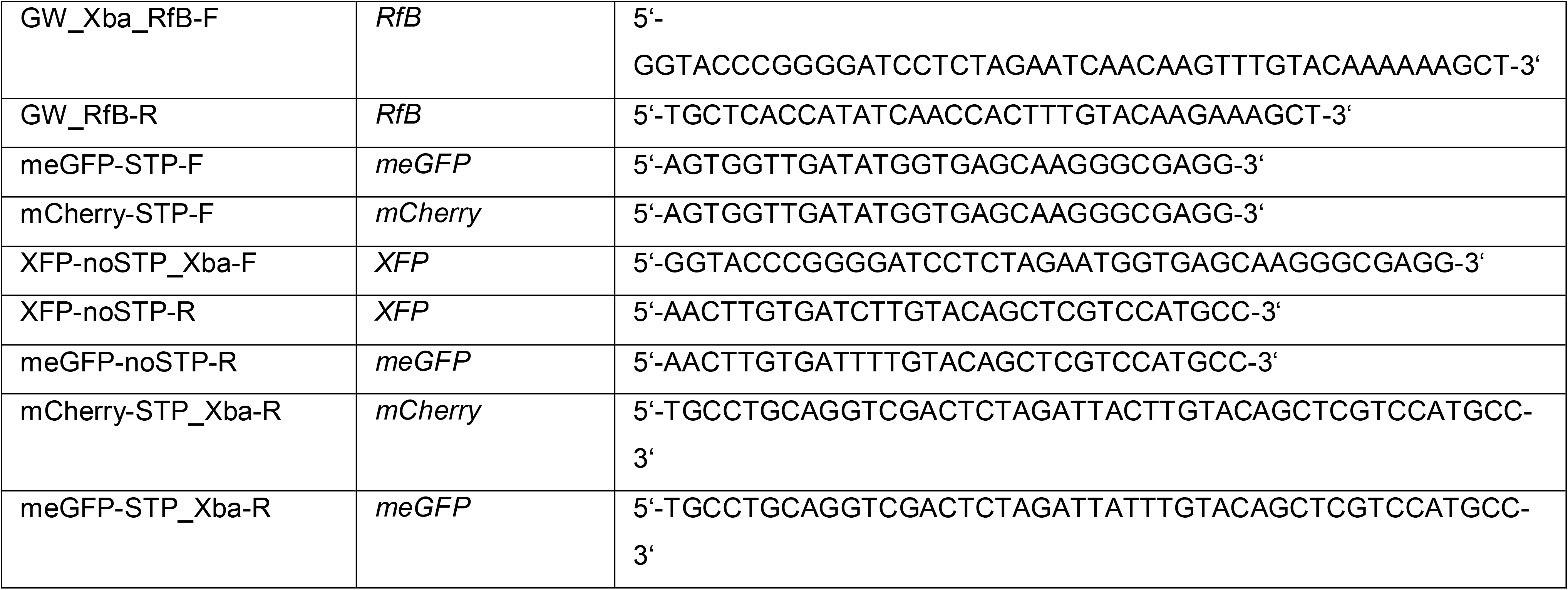
Primers used in this study.

